# Low immunogenicity of malaria pre-erythrocytic stages can be overcome by vaccination

**DOI:** 10.1101/2020.09.03.281238

**Authors:** Katja Müller, Matthew P. Gibbins, Arturo Reyes-Sandoval, Adrian V. S. Hill, Simon J. Draper, Kai Matuschewski, Olivier Silvie, Julius Clemence R. Hafalla

**Affiliations:** Parasitology Unit, Max Planck Institute for Infection Biology, Berlin, Germany; Molecular Parasitology, Institute of Biology, Humboldt University, Berlin, Germany; Department of Immunology and Infection, Faculty of Infectious and Tropical Diseases, London School of Hygiene and Tropical Medicine, London, United Kingdom; Jenner Institute, University of Oxford, Old Road Campus Research Building, Oxford, United Kingdom; Sorbonne Université, INSERM, CNRS, Centre d’Immunologie et des Maladies Infectieuses, CIMI-Paris, Paris, France

**Keywords:** malaria, *Plasmodium*, antigen, vaccine, immunogenicity, protective efficacy, sporozoite, liver stage

## Abstract

Vaccine discovery and development critically depends on predictive assays, which prioritise protective antigens. Immunogenicity is considered one important criterion for progression of candidate vaccines to further clinical evaluation, including phase I/II trials. Here, we tested this assumption in an infection and vaccination model for malaria pre-erythrocytic stages. We engineered *Plasmodium berghei* parasites that harbour a well-characterised epitope for stimulation of CD8+ T cells either as an antigen in the circumsporozoite protein (CSP), the surface coat protein of extracellular sporozoites or in the upregulated in sporozoites 4 (UIS4), a major protein associated with the parasitophorous vacuole membrane (PVM) that surrounds the intracellular exo-erythrocytic forms (EEF). We show that the antigen origin results in profound differences in immunogenicity with a sporozoite antigen eliciting robust and superior antigen-specific CD8+ T cell responses, whilst an EEF antigen evokes poor responses. However, despite their contrasting immunogenic properties, both sporozoite and EEF antigens gain access to antigen presentation pathways in hepatocytes, as recognition and targeting by vaccine-induced, antigen-specific effector CD8+ T cells results in high levels of protection when targeting both antigens. Our study is the first demonstration that poor immunogenicity of EEF antigens does not preclude their susceptibility to antigen-specific CD8+ T cell killing. Our findings that antigen immunogenicity is an inadequate predictor of vaccine susceptibility have wide-ranging implications on antigen prioritisation for the design and testing of next-generation pre-erythrocytic malaria vaccines.

## INTRODUCTION

Malaria, caused by the apicomplexan parasites *Plasmodium*, is responsible for 228 million clinical cases and 405,000 deaths annually worldwide^1^. Whilst current malaria control strategies have led to marked reduction in incidence rate, cases, and mortality for the past 16 years, a highly efficacious vaccine is likely essential to approach the ambitious World Health Organisation’s (WHO) vision of “a world free of malaria”. Targeting the malaria pre-erythrocytic stages, an obligatory and clinically silent phase of the parasite’s life cycle, is considered an ideal and attractive strategy for vaccination; inhibiting parasite infection of and development in hepatocytes results in preclusion of both disease-causing blood stages and transmissible sexual stages. Yet, despite intensive research for over 25 years, a highly efficacious pre-erythrocytic stage vaccine remains elusive^2^. An in-depth characterisation of how the complex biology of pre-erythrocytic stages influences the generation and protective efficacy of immune responses is warranted to inform the design of future malaria vaccines.

CD8+ T cells are crucial mediators of protective immunity to malaria pre-erythrocytic stages^3^. Whilst often considered as a single phase of the parasite’s life cycle, the malaria pre-erythrocytic stage is comprised of two different parasite forms: (i) sporozoites, which are motile extracellular parasites that are delivered by infected mosquitoes to the mammalian host, and (ii) exo-erythrocytic forms (EEF; also known as liver stages), which are intracellular parasites resulting from the differentiation and growth of sporozoites inside a parasitophorous vacuole (PV) within hepatocytes^4^. How these two spatially different parasite forms and the ensuing temporal expression of parasite-derived antigens impact the magnitudes, kinetics and phenotypes of CD8+ T cell responses elicited following infection is poorly understood. Furthermore, the complexity within the pre-erythrocytic stages has fuelled a long-standing debate focused on the contributions of distinct sporozoite and EEF antigens in parasite-induced responses, and whether sporozoite or EEF proteins are better targets of vaccines.

Our current understanding of CD8+ T cell responses to malaria pre-erythrocytic stages has been largely based on measuring responses to the H-2-K^d^-restricted epitopes of *P. yoelii* (*Py*)^5^ and *P. berghei* (*Pb*)^6^ circumsporozoite proteins (CSP), the major surface antigen of sporozoites. Many of these fundamental studies have focused on using infections with irradiated sporozoites, the gold-standard vaccine model for malaria. Infection with *Py* sporozoites elicits an expected T cell response typified by early activation and induction of effector CSP-specific CD8+ T cells followed by contraction and establishment of quantifiable memory populations^7^. CSP-specific CD8+ T cells are primed by dendritic cells that cross-present sporozoite antigens via the endosome-to-cytosol pathway^8^. Yet, CSP is a unique antigen because it is expressed in both sporozoites and EEFs^9^. Whilst the expression of CSP mRNA ceases after sporozoite invasion, the protein on the parasite surface is stable and endures in EEFs during development in hepatocytes^10^. *In vitro* data indicate that primary hepatocytes process and present *Pb*CSP-derived peptides to CD8+ T cells in a proteasome-dependent manner, involving export of antigen to the cytosol^8^. Taken together, these data imply that sporozoite antigens induce quantifiable CD8+ T cell responses after infection. Antigens that have similar expression to the CSP, persisting to EEFs and with epitope determinants presented on hepatocytes, are excellent targets of CD8+ T cell-based vaccines.

The paucity of EEF only-specific epitopes has hindered not only our ability to understand the immune responses that are evoked whilst the parasite is in the liver, but also their utility as targets of vaccination. Accordingly, the contribution of EEF-infected hepatocytes in the *in vivo* induction of CD8+ T cell responses is poorly understood. The liver is an organ where the primary activation of CD8+ T cells is generally biased towards the induction of tolerance^11,12^. Yet, studies in other model systems have demonstrated antigen-specific primary activation within the liver^13^. Another confounding issue with EEFs is their development in PVs with constrained access to the hepatocyte’s cytosol^4^. Nonetheless, if CD8+ T cells specific for EEF antigens are primed, do they expand and contract with distinct kinetics? Moreover, are EEF-specific epitopes efficiently generated for recognition and targeting by vaccine-induced CD8+ T cells? Answers to these questions will be key for antigen selection and design of future malaria vaccines.

In this study, we compared the initiation and development of CD8+ T cell responses – elicited following parasite infection – to CSP, a sporozoite antigen, and to upregulated in infective sporozoites gene 4 (UIS4), an EEF-specific vacuolar protein^14^. UIS4, a member of the early transcribed membrane protein (ETRAMP) family, is abundantly expressed in EEFs and associates with the PVM^14^. Whilst UIS4 mRNA expression is present in sporozoites, translation is repressed until when EEFs develop^10^. To control for epitope specificity, we generated *Pb* transgenic parasites that incorporate the H-2-K^b^ epitope SIINFEKL, from ovalbumin, in either CSP or UIS4. Furthermore, we evaluated the capacity of vaccine-induced CD8+ T cells to target these parasites in a mouse challenge model. Our data show disparate immunogenic properties between a sporozoite and an EEF vacuolar membrane antigen but equivalent susceptibility to vaccine-induced CD8+ T cells.

## RESULTS

### Transgenic CSP^SIINFKEL^ and UIS4^SIINFEKL^ parasites display normal sporozoite motility and liver invasion

We generated, by double homologous recombination, transgenic *Pb* parasites expressing the immunodominant H-2-K^b^-restricted CD8+ T cell epitope of ovalbumin (SIINFEKL) in the context of the sporozoite surface antigen CSP or the EEF vacuolar membrane antigen UIS4 **(Figure 1a and Supplementary Figure 1a, b)**. Constructs included the *TgDHFR/TS* positive selection cassette and incorporated SIINFEKL in the context of the gene open reading frame. For CSP^SIINFEKL^, SIINFEKL replaced SYIPSAEKI, the immunodominant H-2-K^d^-restricted CD8+ T cell epitope of CSP, which allowed for recognition in H-2-K^b^-carrying C57BL/6 mice. For UIS4^SIINFEKL^, the SIINFEKL epitope was added to the immediate C-terminus of the UIS4 protein. Appending the C-terminus was chosen because it had been shown in *Toxoplasma gondii* that the potency of the immunodominant epitope of GRA6 was associated with its C-terminal location, which may have enhanced the presentation by parasite-infected cells^15^. Whilst undefined for UIS4 itself, it has been shown for several other ETRAMPs that the C-terminus faces the host cell cytoplasm^16^, which might enhance exposure to the MHC I machinery.

**Figure 1:**
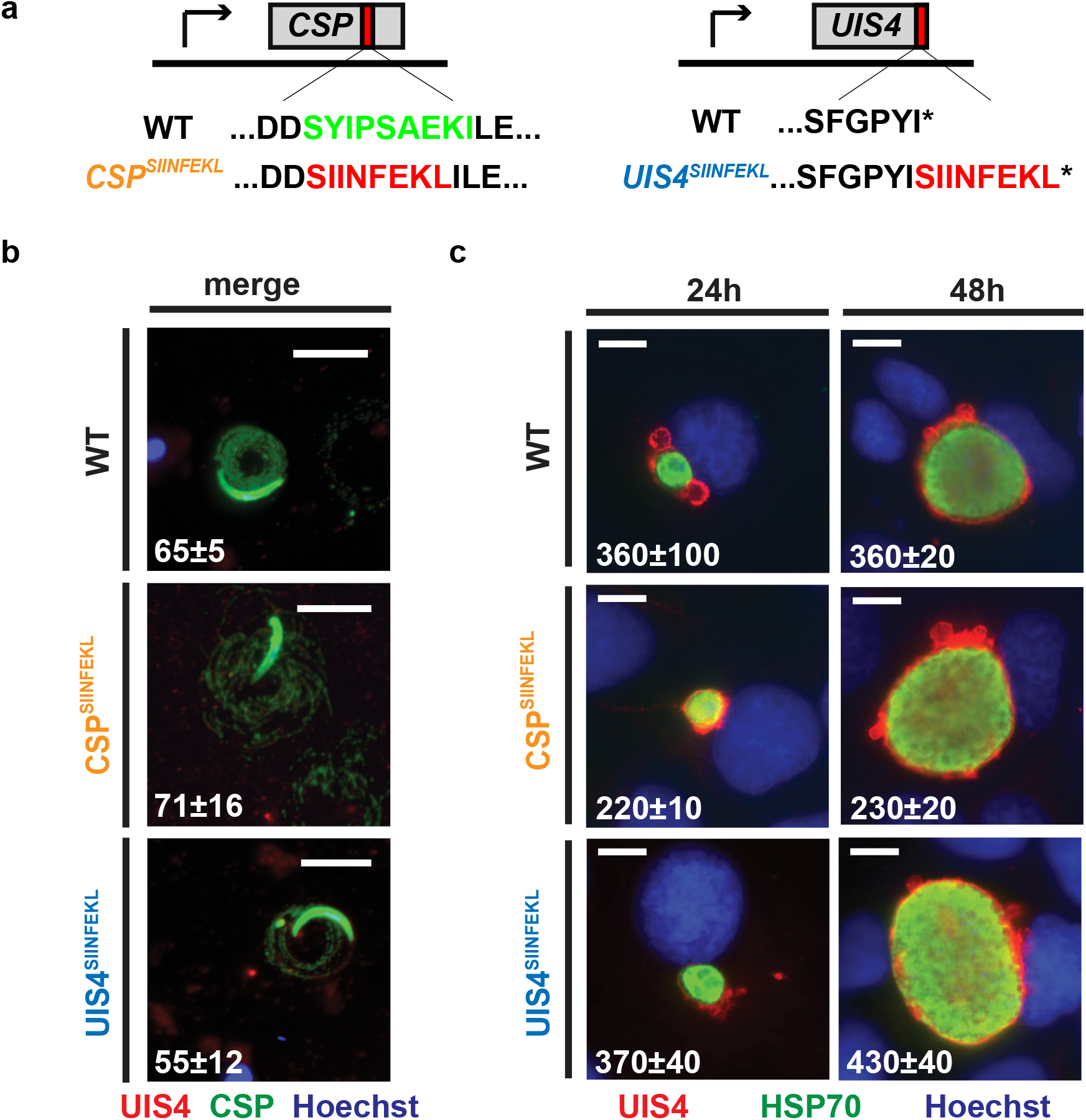
Generation and characterisation of recombinant CSP^SIINFEKL^ and UIS4^SIINFEKL^ *P. berghei* parasites. *Pb* parasites expressing the CD8+ T cell epitope of ovalbumin, SIINFEKL, in the context of CSP or UIS4 were generated using double homologous recombination. **(a)** To generate CSP^SIINFEKL^, SIINFEKL replaced amino acids SYIPSAEK in CSP. To generate UIS4^SIINFEKL^, SIINFEKL was adjoined to the carboxyl-terminus of the UIS4 protein. **(b)** Sporozoite immunofluorescent antibody staining of WT, CSP^SIINFEKL^ or UIS4^SIINFEKL^ sporozoites after gliding on BSA-coated glass slides. Shown are microscopic images of the respective sporozoites that were stained with anti-CSP (green), anti-UIS4 (red) and nuclear stain Hoechst 33342 (blue). Scale bars, 10μm. The numbers show mean percentage (±SD) of sporozoites with trails assessed from ≥220 sporozoites. **(c)** Fluorescent-microscopic images of EEF-infected Huh7 hepatoma cells. 24 and 48 hours after infection with WT, CSP^SIINFEKL^ or UIS4^SIINFEKL^ sporozoites, the cells were fixed and stained with anti-UIS4 (red), anti-HSP70 (green) and the nuclear stain Hoechst (blue). Scale bars: 10μm. The numbers show mean numbers (±SD) of intracellular parasites counted from ≥200 EEFs.

The resulting parasites showed a phenotype comparable to WT parasites, with functional sporozoite motility **(Figure 1b)** and normal invasive capacity and development inside hepatocytes **(Figure 1c)**, as well as comparable midgut infectivity and number of salivary gland sporozoites **(Supplementary Figure 1c, d)**. Thus, the introduced mutations to generate CSP^SIINFEKL^ and UIS4^SIINFEKL^ parasites did not interfere with the completion of the life cycle, in either mosquito vector or mouse. All C57BL/6 mice that received 800 sporozoites of either CSP^SIINFKEL^ or UIS4^SIINFEKL^ intravenously developed a patent blood stage infection by day 4, comparable to infection with WT sporozoites (data not shown).

### Peripheral blood CD8+ T cell responses and early proliferative capacity of splenic CD8+ T cells are superior if elicited by a sporozoite surface protein in contrast to a vacuolar membrane protein in the infected liver

We first wanted to determine whether the generated transgenic parasites allow antigen-specific responses to be tracked using SIINFEKL as a surrogate CD8+ T cell epitope for sporozoite surface and EEF vacuolar membrane antigens. To this end, we assessed the kinetics of the CD8+ T cell response following intravenous immunisation with CSP^SIINFEKL^ or UIS4^SIINFEKL^ sporozoites. To augment the CD8+ T cell response, mice were adoptively transferred with 2 × 10^6^ OT-I cells expressing a SIINFEKL-specific TCR^8^, prior to receiving 10,000 γ-radiation attenuated WT, CSP^SIINFEKL^ or UIS4^SIINFEKL^ sporozoites. Prior work showed that γ-radiation attenuation of *P. berghei* sporozoites does not impact host cell invasion and UIS4 expression^17^.

Peripheral blood was taken at days 4, 7, 14, 21, 42 and 88 after immunisation and CD8+ T cell responses were analysed after staining with H-2-K^b^-SIINFEKL pentamers and for CD11a, a marker for antigen-experienced T cells^18,19^ (**Figure 2a)**. A substantial proportion of K^b^-SIINFEKL+ CD11a+ CD8+ T cells were observed in mice immunised with CSP^SIINFEKL^; the response was highest on day 4, reaching 5% of all antigen-experienced CD8+ T cells, and declined steadily until day 21, when the response stabilised and remained unchanged for several weeks **(Figure 2b)**. In marked contrast, UIS4^SIINFEKL^ immunisation induced a poor CD8+ T cell response; the proportion of K^b^-SIINFEKL+ CD11a+ CD8+ T cells was only higher than the control groups at day 4 after immunisation, and the response remained within background levels for the duration of the experiment. Control groups included mice receiving OT-I cells only or in addition to WT sporozoites, which lack SIINFEKL sequences.

**Figure 2:**
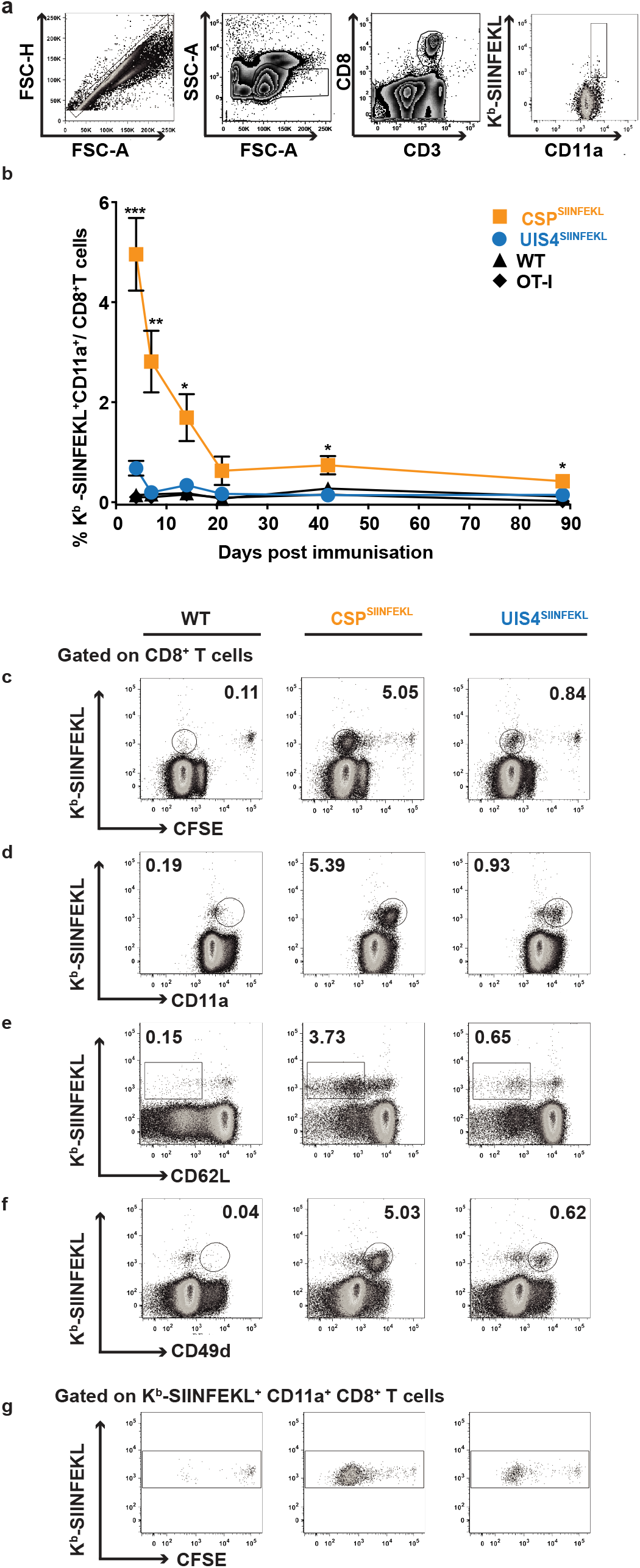
Kinetics of CD8+ T cell responses induced by transgenic parasites. **(a-b)** C57BL/6 mice (up to 8 per group) received 2×10^6^ OT-I cells alone (diamonds) or were additionally immunised with 10,000 γ-radiation attenuated WT (triangles), CSP^SIINFEKL^ (orange squares) or UIS4^SIINFEKL^ (blue circles) sporozoites intravenously. **(a)** Flow cytometry plots show the gating strategy for identifying K^b^-SIINFEKL+ CD11a+ CD8+ T cells. **(b)** Peripheral blood was obtained on days 4, 7, 14, 21, 42 and 88 after immunisation and stained for K^b^-SIINFEKL+ CD11a+ CD8+ T cells. Line graph shows mean values (±SEM) from representative experiments (*, p<0.05; **, p<0.01; ***, p<0.001; Welch’s t-test). **(c-g)** C57BL/6 mice (n=4), which received 2×10^6^ CFSE-labelled OT-I splenocytes, were immunised with 10,000 γ-radiation attenuated WT, CSP^SIINFEKL^ or UIS4^SIINFEKL^ sporozoites intravenously. 5 days later, mice were sacrificed, spleens harvested and splenocytes assessed for **(c)** CFSE dilution and stained *ex vivo* **(d-f)** for effector CD8+ T cell surface markers. Shown are flow cytometry plots of K^b^-SIINFEKL co-staining with markers of effector phenotypes: **(d)** CD11a^hi^, **(e)** CD62L^lo^, **(f)** CD49d^hi^ and **(g)** the proliferation of CFSE-labelled antigen experienced Kb-SIINFEKL+ CD11a+ CD8+ T cells.

The poor CD8+ T cell response induced by UIS4^SIINFEKL^ sporozoites, as compared to CSP^SIINFEKL^, led us to characterise the early events in the proliferation and differentiation of these cells. Mice were adoptively transferred with CFSE-labelled OT-I cells and immunised with γ-radiation attenuated WT, CSP^SIINFEKL^ or UIS4^SIINFEKL^ sporozoites. As shown by gating on CD8+ T cells **(Figure 2c, g**, after 5 days, immunisation with CSP^SIINFEKL^ sporozoites recruited K^b^-SIINFEKL+ CD8+ T cells to undergo massive proliferative activity, which was 6x larger than that observed with UIS4^SIINFEKL^ sporozoites, in good agreement with the peripheral blood data described above **(Figure 2b)**. Consistent with the activation of these cells, the proliferation of antigen-specific CD8+ T cells by both parasites was associated with the development of effector and effector-memory phenotypes as evidenced by upregulation of CD11a and CD49d, and downregulation of CD62L, respectively **(Figure 2d-f)**.

Taken together, these findings establish that immunisations with CSP^SIINFEKL^ and UIS4^SIINFEKL^ sporozoites permit antigen-specific responses to be tracked longitudinally in the peripheral blood. Importantly, we demonstrate that a sporozoite surface protein evokes a CD8+ T cell response of superior magnitude than an EEF vacuolar membrane protein following immunisation with malaria sporozoites.

### High magnitude splenic and hepatic CD8+ T cell responses to a sporozoite antigen

Previous research has shown that CD8+ T cells are primed primarily in the spleen following intravenous immunisation with malaria sporozoites^20^ and that liver lymphocytes form a front-line defence against developing EEFs in hepatocytes^21,22^. Thus we further analysed the development of CD8+ T cell responses in the spleens and livers of mice adoptively transferred with OT-I cells and intravenously immunised with WT, CSP^SIINFEKL^ or UIS4^SIINFEKL^ sporozoites. Consistent with our aforementioned results, surface staining of splenic and liver lymphocytes showed higher proportion and absolute numbers of K^b^-SIINFEKL+ CD11a+ CD8+ T cells at day 14 and day 42 following immunisation with CSP^SIINFEKL^ compared to UIS4^SIINFEKL^ sporozoites **(Figure 3a-c)**. In addition to CD11a upregulation, the splenic and liver CD8+ T cells, elicited by both CSP^SIINFEKL^ or UIS4^SIINFEKL^ sporozoites, had effector and effector memory cell phenotypes (CD62L-, CD49d+ and CD44+) **(Supplementary Figure 2)**. Although low, the numbers of antigen-specific CD8+ T cells induced by UIS4^SIINFEKL^ sporozoites were within the detection limits of the assay.

**Figure 3:**
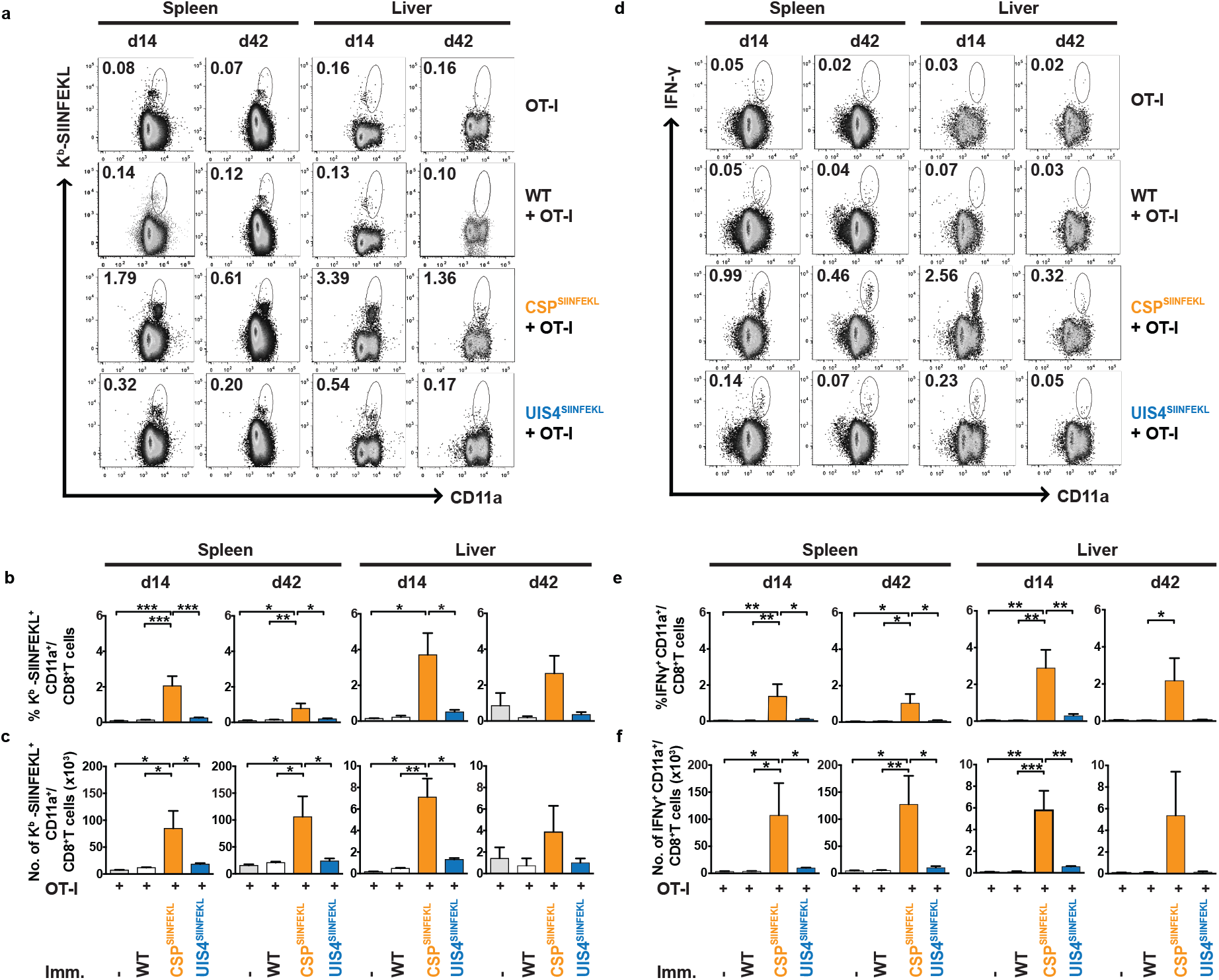
Sporozoite surface antigen induces a higher CD8+ T cell response than EEF vacuolar membrane antigen in the spleen and liver. C57BL/6 mice (up to 5) received 2×10^6^ OT-I cells alone or were additionally immunised with 10,000 γ-radiation attenuated WT, CSP^SIINFEKL^ or UIS4^SIINFEKL^ sporozoites intravenously. Spleens and livers were harvested either at day 14 or day 42. Proportions and numbers of **(a-c)** K^b^-SIINFEKL+ CD8+ T cells were enumerated or **(d-f)** IFN-γ-secreting CD8+ T cells following restimulation *ex vivo* with SIINFEKL peptide were quantified. Flow cytometry plots show representative percentages of CD8+ T cells co-stained with CD11a and **(a)** K^b^-SIINFEKL or **(d)** IFN-γ. The upper panel of bar charts **(b, e)** show the percentage of co-stained CD8+ T cells and the lower panel **(c, f)** the absolute cell counts. Bar charts show mean values (±SEM) from representative experiments (*, p<0.05; **, p<0.01; ***, p<0.001; one-way ANOVA with Tukey’s multiple comparison test).

To assess for effector functions, splenic and liver lymphocytes were stimulated *ex vivo* with the SIINFEKL peptide. Generally, higher numbers (proportion and absolute numbers) of IFN-γ-secreting CD8+ T cells were observed at day 14 and day 42 following immunization with CSP^SIINFEKL^ compared to UIS4^SIINFEKL^ sporozoites **(Figure 3d-f)**. In addition, these CD8+ T cells also expressed TNF and IL-2, suggesting some potential polyfunctionality **(Supplementary Figure 3)**.

Altogether, even though effector and effector memory CD8+ T cell responses can be detected against both sporozoite surface protein and EEF vacuolar membrane protein antigens following immunisation with malaria sporozoites, the two antigens show a striking difference in the magnitude of CD8+ T cell responses they induce.

### Quantification of endogenously produced antigen-specific CD8+ T cells following intravenous or intradermal parasite immunisation

Previous work tracking responses to SIINFEKL-tagged proteins has used adoptively transferred cells from OT-I mice, with all T cells from these mice expressing T cell receptors specific to SIINFEKL^8,23^. We employed this robust approach by adoptively transferring a fixed amount of OT-I splenocytes in order to augment the response and allow visualisation **(Figures 2 and 3)**. Next, we wanted to explore whether we can capture the endogenous K^b^-SIINFEKL+ CD11a+ CD8+ T cell population, which is elicited by immunising with parasites without OT-I cell transfer. We performed *ex vivo* restimulation of lymphocytes with SIINFEKL peptide followed by flow cytometry and were able to clearly identify the endogenous population with a trend complementary to our earlier results **(Figure 4a-c)**. Immunisation with CSP^SIINFEKL^ sporozoites elicited a superior splenic and liver CD8+ T cell response than with UIS4^SIINFEKL^ sporozoites. As expected, the proportion and absolute cell numbers were considerably lower than with adoptive transfer of OT-I cells, but this did not preclude the ability to visualise IFN-γ-secreting CD8+ T cells and capture the differences between the two groups.

**Figure 4:**
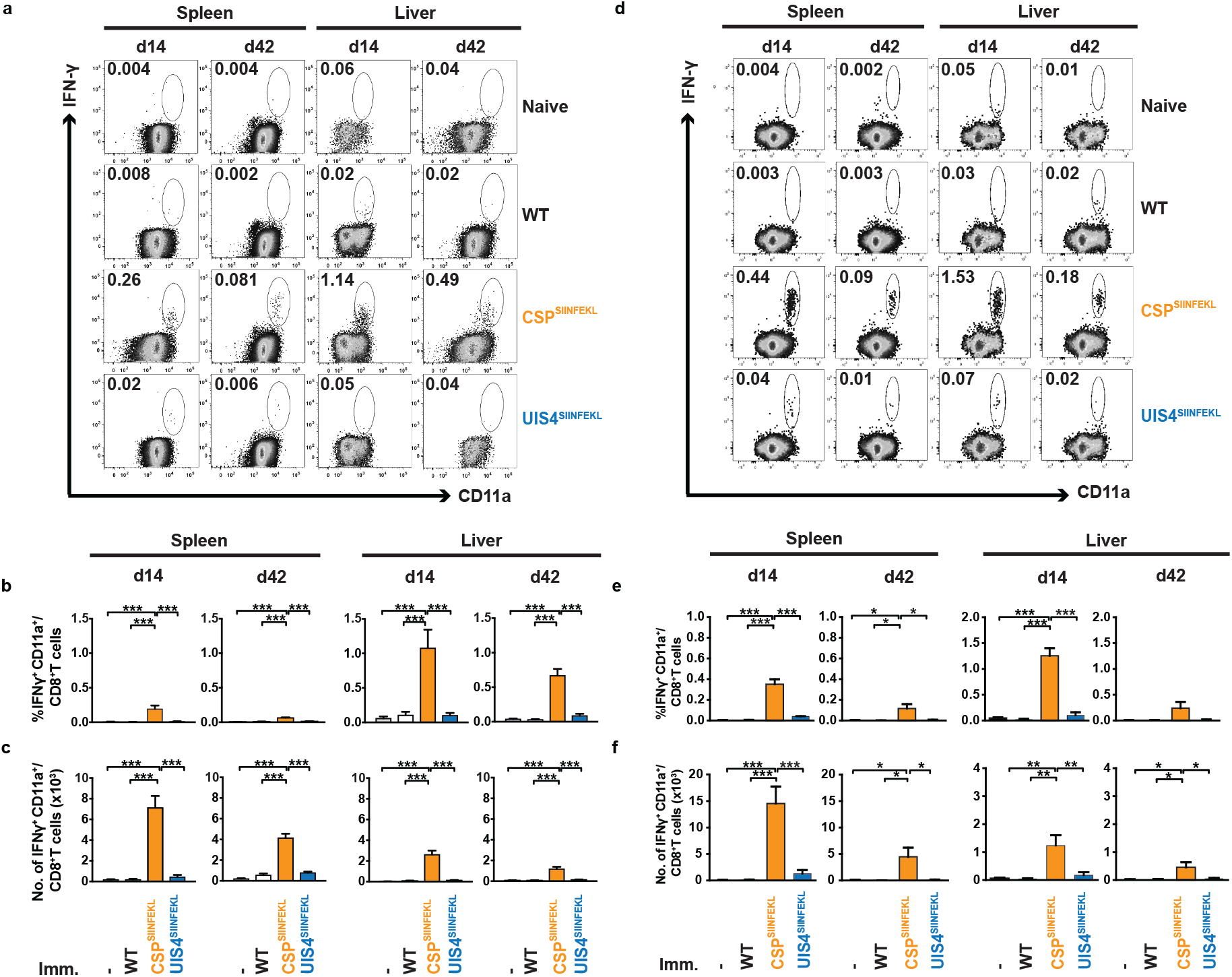
OT-I cells are not required to detect SIINFEKL-specific CD8+ T cell responses. C57BL/6 mice (3-6 per group) received 10,000 γ-radiation attenuated WT, CSP^SIINFEKL^ or UIS4^SIINFEKL^ sporozoites, either **(a-c)** intravenously or **(d-f)** intradermally. Additional control mice did not receive sporozoites. Spleens and livers were harvested either at day 14 or day 42, and IFN-γ-secreting lymphocytes following restimulation *ex vivo* with SIINFEKL peptide were quantified. Flow cytometry plots show representative percentages of CD8+ T cells co-stained with IFN-γ and CD11a **(a, d)**. The upper panel of bar charts **(b, e)** show the percentage of CD11a+ IFN-γ+ CD8+ T cells and the lower panel **(c, f)** the absolute cell counts. Bar charts show mean values (±SEM) from representative experiments (*, p<0.05; **, p<0.01; ***, p<0.001; one-way ANOVA with Tukey’s multiple comparison test).

Under normal conditions of transmission, sporozoites are delivered into the host skin by mosquito bite. All preceding immunisation experiments were performed with parasites injected intravenously. As a proxy for the natural route of infection, whilst ensuring consistent quantities of parasites were inoculated, CSP^SIINFEKL^ and UIS4^SIINFEKL^ sporozoites were injected via the intradermal route into the ear pinnae. Under these conditions, CSP still induced a greater number of IFN-γ-secreting SIINFEKL-specific CD8+ T cells following restimulation with SIINFEKL compared to UIS4, with a comparable magnitude as after intravenous injection **(Figure 4d-f)**. Thus, these biologically and immunologically more appropriate data entirely recapitulate the strong immunogenicity of a sporozoite surface antigen compared to an EEF vacuolar membrane protein.

### Increasing the amount of EEF vacuolar membrane antigen does not impact its immunogenicity

Both CSP and UIS4 are critical proteins expressed by the sporozoite and EEF respectively, and both proteins are important for survival and succession into the subsequent life stage and parasite form^10,14,24^. Previous studies have shown that the magnitude of the CD8+ T cell response to a sporozoite surface antigen depended on the amount of parasites used for immunisation^25^. Hence, poor immunogenicity of an EEF vacuolar membrane protein could be a result of the lower level of protein expression during parasite infection. It is possible to enhance CD8+ T cell responses by increasing the number of parasites used for immunisation^25^. Therefore, we immunised groups of mice with 8,000 CSP^SIINFEKL^, 8,000 UIS4^SIINFEKL^ or 64,000 UIS4^SIINFEKL^ sporozoites and compared the magnitude of the elicited antigen-specific responses. Strikingly, the CD8+ T cell response following 8x sporozoite immunisation dose with UIS4^SIINFEKL^ did not increase proportionally and was not significantly higher than immunisation with a 1x dose **(Figure 5a, b)**. This result suggests that, in the context of attenuated sporozoite immunisation, EEF vacuolar membrane antigens are poorly immunogenic and increasing antigen fails to substantially improve the magnitude of CD8+ T cell responses.

**Figure 5:**
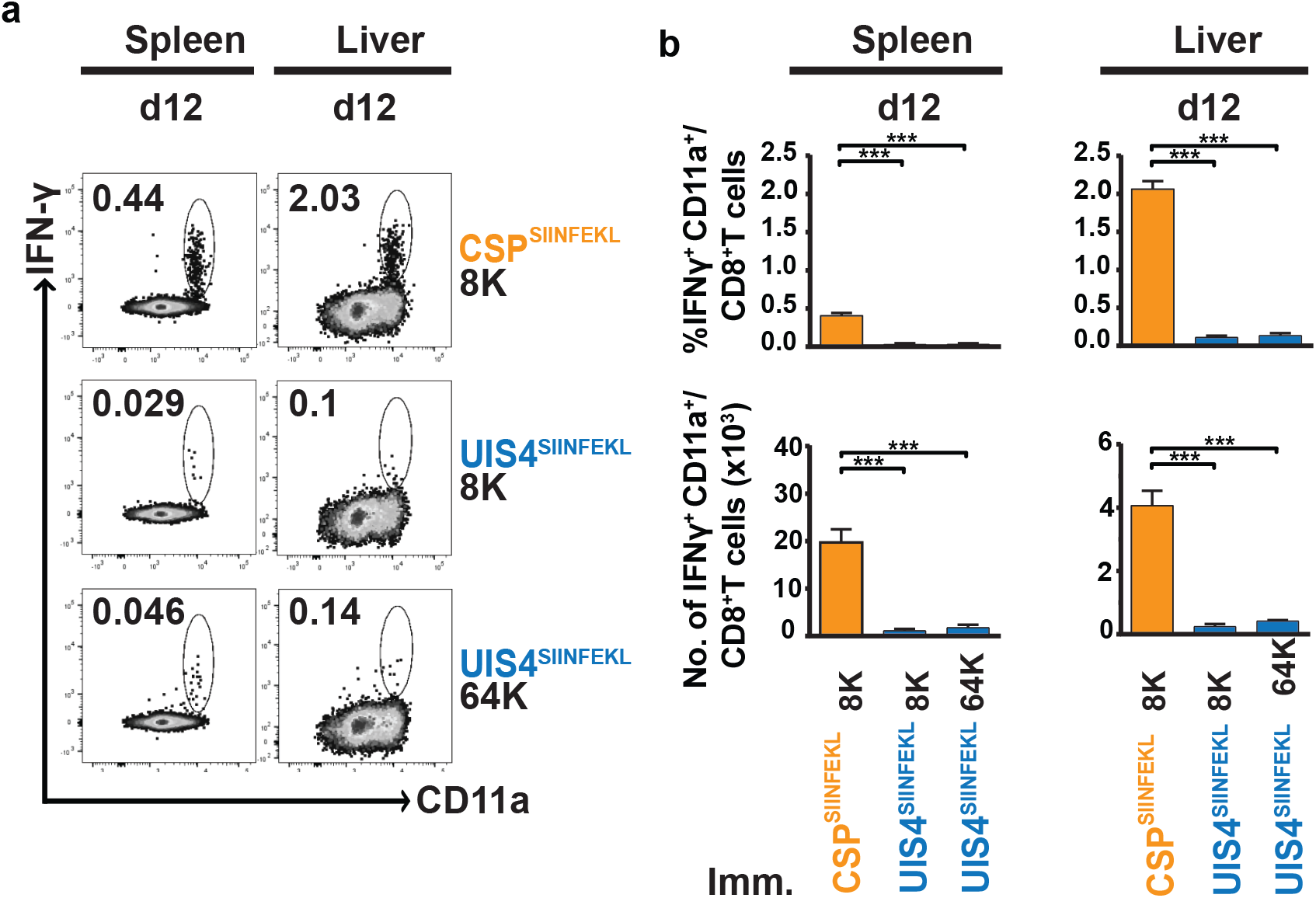
Increasing antigen dose does not improve antigen-specific CD8+ T cell responses to an EEF vacuolar membrane protein. C57BL/6 mice (4 per group) received an intravenous dose of 8,000 γ-radiation attenuated CSP^SIINFEKL^ or UIS4^SIINFEKL^ sporozoites or 64,000 γ-radiation attenuated UIS4^SIINFEKL^ sporozoites. Spleens and livers were harvested at day 12 and IFN-γ-secreting lymphocytes following restimulation *ex vivo* with SIINFEKL peptide were quantified. **(a)** Flow cytometry plots show representative CD8+ T cells co-stained with IFN-γ and CD11a. **(b)** The upper panel of bar charts show the percentage of CD11a+ IFN-γ+ CD8+ T cells and the lower panel the absolute cell counts. Bar charts show mean values (±SEM) from representative experiments (***, p<0.001; one-way ANOVA with Tukey’s multiple comparison test).

### Immunogenicity of parasite antigens does not predict effector responses following vaccination

Our findings thus far showed that sporozoite surface proteins appear more immunogenic than EEF vacuolar membrane proteins and raised an intriguing and important question; does immunogenicity predict susceptibility to vaccine-induced effector responses? To address this, we vaccinated mice, which had received OT-I cells, with a recombinant adenovirus expressing full-length ovalbumin^26^. This vaccination protocol resulted in frequencies of ~7.5% SIINFEKL-specific CD8+ T cells in peripheral blood **(Figure 6a, b)**. Vaccinated mice were then challenged with CSP^SIINFEKL^ or UIS4^SIINFEKL^ sporozoites, and protection was assessed 19 days after vaccination by two complementary assays; (i) determination of the reduction of parasite load in the liver 42 hours after sporozoite challenge **(Figure 6c)**, and (ii) induction of sterile protection **(Figure 6d)**. Vaccinated mice challenged with CSP^SIINFEKL^ or UIS4^SIINFEKL^ sporozoites showed a dramatic reduction in parasite load in the liver **(Figure 6c)** as compared to vaccinated mice challenged with WT parasites. Strikingly, there was no statistical difference in the protection observed when vaccinated mice were challenged with either CSP^SIINFEKL^ or UIS4^SIINFEKL^ sporozoites. Consistent with these findings, both groups of vaccinated mice challenged with either CSP^SIINFEKL^ or UIS4^SIINFEKL^ sporozoites exhibited sterile protection of comparable levels **(Figure 6d)**. These findings indicate that spatial and temporal aspects of antigen expression may affect protein immunogenicity in the context of parasitic infection but not necessarily the same target’s susceptibility for antigen-specific CD8+ T cell killing.

**Figure 6:**
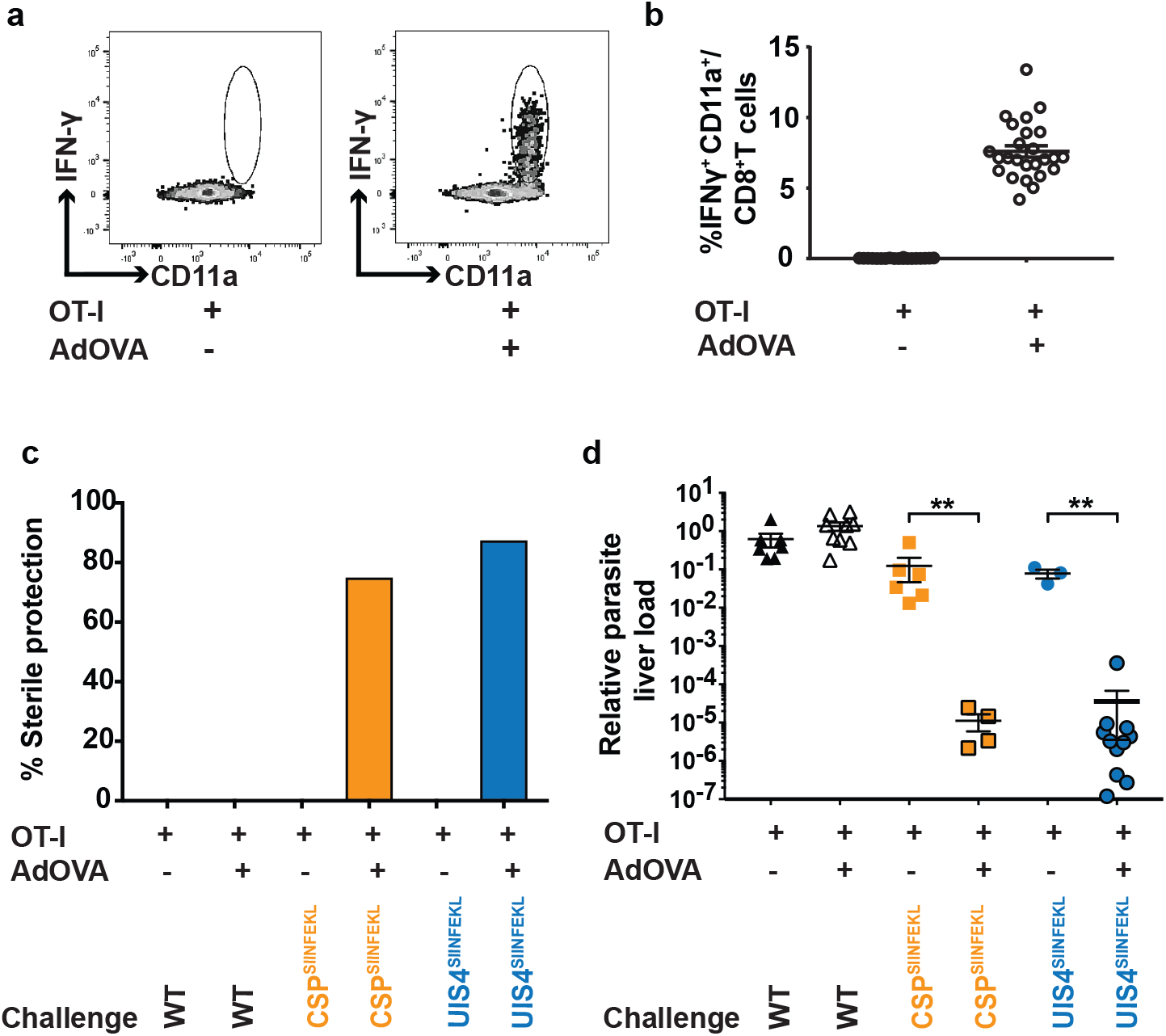
Sporozoite surface and EEF vacuolar membrane antigens are presented to vaccine-induced CD8+ T cells for killing, leading to sterile protection. Mice received 1×10^8^ ifu recombinant AdHu5 expressing whole ovalbumin (AdOVA) and/or 2×10^6^ OT-I splenocytes. **(a)** Flow cytometry and **(b)** scatter plots represent CD8+ T cells derived from peripheral blood co-stained with IFN-γ and CD11a, following *ex vivo* restimulation with SIINFEKL. **(c)** Protective efficacy as measured by quantitative real-time PCR. Groups of mice (up to 11 per group) were vaccinated as described and challenged 19 days later with 10,000 WT, CSP^SIINFEKL^ or UIS4^SIINFEKL^ sporozoites. 42 hours later livers were removed and parasite load was assessed by qPCR. Plots show the relative parasite load of mice in each condition (**, p<0.01; Mann-Whitney U test). **(d)** Proportion of sterile protection after immunization. Mice (8 per group) were vaccinated as described and were challenged with 1,000 WT, CSP^SIINFEKL^ or UIS4^SIINFEKL^ sporozoites. Data for **a-d** are representative of two experiments performed with scatter plots showing mean values (±SEM).

## DISCUSSION

The malaria pre-erythrocytic stages have been a prime target for the development of a *Pf* vaccine for more than 35 years. Indeed, RTS,S/AS01, the most advanced malaria sub-unit vaccine candidate to date is based on CSP, the major surface protein of sporozoites^27^. Yet, final results of the Phase III trial showed that RTS,S/AS01 offers only modest efficacy, which rapidly wanes over time^28^. Thus, there is an imperative need not only to widen the pursuit for new sub-unit vaccine candidates, but also to radically improve the antigen selection process. Antigens are generally prioritised based on a range of criteria, including their immunogenicity in the context of parasitic infection. We examined this notion in an infection and vaccination model for malaria pre-erythrocytic stages.

The malaria pre-erythrocytic stages consist of two spatially-different parasite forms: extracellular sporozoites and intracellular EEFs. The transformation of sporozoites into EEFs involves regulation at both transcriptional^29^ and translational^30,31^ levels, resulting in both the spatial and temporal expression of many antigens that are distinct for each parasite form^32^. Whilst our current understanding of immune responses to malaria pre-erythrocytic stages has focused on CSP, the lack of well-defined epitopes that are expressed only by EEFs has restrained fundamental studies investigating the contributions of EEF antigens in parasite-induced CD8+ T cell responses and their value as target of vaccines.

In this study, we contrasted the development of CD8+ T cell responses induced by CSP and UIS4, two major proteins expressed by sporozoites and EEFs, respectively. We generated transgenic *Pb* parasites where SIINFEKL is expressed as part of either CSP or UIS4, allowing the presentation of the epitope at the same space and time as the respective protein. This approach is in contrast to a more common strategy of expressing the whole, or a fragment of, ovalbumin inserted as a transgene into the *Pb* genome, and then tracking the immune response elicited by an extraneous molecule^23,33^. Since CSP is expressed in both sporozoites and EEFs, the processing and presentation of the SIINFEKL in CSP^SIINFEKL^ occurs as soon as sporozoites are inoculated and are able to interact with dendritic cells, which present antigens via an endosome-to-cytosol pathway^8^; CSP also has direct access to the hepatocyte’s cytosol for processing and presentation of the CSP-derived epitope^8^. However, since UIS4 is expressed only in the PVM of EEFs, processing and presentation of the epitope in UIS4^SIINFEKL^ is restricted to just hepatocytes.

Our results establish that following sporozoite-immunisation, a sporozoite surface protein induces superior CD8+ T cell responses – as measured both by pentamer staining and by IFN-γ secretion following peptide stimulation – than an EEF vacuolar membrane protein. Detailed kinetic and phenotypic analysis of the development of antigen-specific CD8+ T cells to both CSP and UIS4 revealed that the responses differ in magnitude, demonstrating the ability of both antigens to elicit effector and effector memory responses. There was no difference in our results whether sporozoites are delivered using the commonly used intravenous immunisation or the more physiological intradermal delivery. We also showed that increasing the number of UIS4^SIINFEKL^ parasites used for immunisation did not augment CD8+ T cell responses, signifying that the poor immunogenicity of an EEF vacuolar membrane protein is not due to the level of UIS4 expression during parasite infection. Our findings support the idea that EEF antigens have minimal contributions to the magnitude of immune responses following whole sporozoite immunisation, which corroborates with prior data showing that that hepatocytes are poor at priming T cell responses^11,12^.

Regardless of their differing immunogenicities in the context of parasitic infection, we further demonstrated that both sporozoite and EEF antigens are effectively targeted by antigen-specific effector CD8+ T cells, which were generated by vaccination using priming and boosting with recombinant viruses expressing the epitope. Importantly, mice harbouring vaccine-induced, antigen-specific CD8+ T cells were comparably protected when challenged with either CSP^SIINFEKL^ or UIS4^SIINFEKL^. These findings imply that both sporozoite and EEF antigens comparably access the antigen presentation pathways in hepatocytes leading to recognition of defined epitopes.

Our study is the first demonstration that poor natural immunogenicity, in this case of an EEF antigen, does not preclude antigen-specific CD8+ T cell killing. Our findings that antigen immunogenicity in this context is an inadequate predictor of vaccine efficacy have wide-ranging implications on antigen prioritisation for the design and testing of next-generation pre-erythrocytic malaria vaccines. Thus, the strategy to screen for T cell responses in naturally infected or sporozoite-immune volunteers to prioritise vaccine candidates requires some form of reassessment. It is noteworthy that for other stages of malaria infection, antigens that give limited or no responses e.g. RH5^34,35^ and sexual stage antigens^36^, are promising antibody targets for vaccines.

A key direction for future research will be identifying the mechanisms by which EEF antigens elicit protection and finding new assays to easily distinguish good vaccine targets, namely those antigens that can protect (via susceptibility to vaccine-induced CD8+ T cells), rather than those that naturally induce strongly immunogenic responses. Ultimately, the molecular mechanisms of presentation of EEF antigens, those expressed in the PVM and within the parasite itself, onto the surface of infected hepatocytes remains to be fully understood. Determination of the processes involved in parasite antigen presentation in the pre-erythrocytic stages of malaria may elucidate links to protection and the identification of further antigens that could drive the development of an efficacious protective malaria vaccine.

## METHODS

### Ethics and animal experimentation

Animal procedures were performed in accordance with the German ‘Tierschutzgesetz in der Fassung vom 18. Mai 2006 (BGBl. I S. 1207)’ which implements the directive 2010/6 3/EU from the European Union. Animal experiments at London School of Hygiene and Tropical Medicine were conducted under license from the United Kingdom Home Office under the Animals (Scientific Procedures) Act 1986. NMRI, CD-1, C57BL/6 and OT-I laboratory mouse strains were bred in house at LSHTM or purchased from Charles River Laboratories (Margate, UK or Sulzfeld, Germany). Female mice were used for experiments at the age of 6-8 weeks.

### Generation of transgenic parasites

Transgenic *P. berghei* ANKA mutants CSP^SIINFEKL^ and UIS4^SIINFEKL^ were developed using double homologous recombination. In the CSP^SIINFEKL^ mutant, the CSP gene is altered so the epitope SYIPSAEKI (residues 252-260) is replaced with the H-2^b^ restricted *Gallus gallus* ovalbumin epitope SIINFEKL. In the UIS4^SIINFEKL^ mutant, the SIINFEKL epitope is appended to the C-terminal end of the UIS4 protein. Clonal parasite lines were generated by limiting dilution. Details of plasmid design, including the primers used and the cloning of parasites can be found in Supplementary Experimental Procedures and Table S1.

### *Plasmodium berghei ANKA* immunisation

*P. berghei* wild type (WT; strain ANKA clone c15cy1 or clone 507) parasites and CSP^SIINFEKL^ and UIS4^SIINFEKL^ (clone c15cy1) parasites were maintained by continuous cycling between murine hosts (NMRI or CD-1) and *Anopheles stephensi* mosquitoes. Infected mosquitoes were kept in incubators (Panasonic and Mytron) at 80% humidity and 20°C. Sporozoites were isolated from salivary glands and γ-irradiated at 1.2 × 10^4^ cGy. Mice were immunised intravenously in the lateral tail vein or intradermally in the ear pinnae with 10,000 sporozoites, unless otherwise stated, and challenged with either 1,000 or 10,000 sporozoites injected intravenously.

### Indirect fluorescent antibody staining (IFA) of sporozoites

Epoxy-covered 8-well glass slides were coated with 3% BSA-RPMI. 10,000 sporozoites were added per well in 3% BSA-RPMI and incubated for 45 minutes during which the shed surface proteins are deposited in the gliding motility process. Sporozoites and their trails were stained with a mouse anti-CSP^37^ primary antibody and a rabbit polyclonal anti-*Pb*UIS4^30^ primary antibody and the respective fluorescently labelled secondary antibodies. Nuclei were stained with Hoechst 33342 and slides mounted with ‘Fluoromount-G’ (Southern Biotech). Sporozoites and trails were analysed by fluorescent microscopy (Zeiss Axio Observer).

### *In vitro* infection of hepatoma cells and fluorescent staining

*In vitro* EEF development was analysed in infected Huh7 hepatoma cells for 24 and 48 hours. Triplicate Labtek (Permanox plastic - Nunc) wells were infected with 10,000 transgenic CSP^SIINFEKL^ or UIS4^SIINFEKL^ parasites and duplicate wells were infected with 10,000 WT parasites. Infected cells were analysed by fluorescence microscopy using a mouse anti-*Pb*HSP70^38^ and a rabbit polyclonal anti-*Pb*UIS4^30^ primary antibody, the respective fluorescently labelled secondary antibodies and nuclear staining with Hoechst 33342. Stainings were analysed by fluorescent microscopy (Zeiss Axio Observer).

### Quantification of SIINFEKL-specific CD8+ T cell responses

Spleens and livers were harvested from immunised or naïve mice and perfused with PBS. Lymphocytes were derived from spleens by passing through 40 or 70μm cell strainers (Corning) and from livers by passing through 70μm cell strainers (Corning). Red blood cells were lysed with PharmLyse (BD), and lymphocytes were resuspended in complete RPMI (cRPMI- RPMI + 10% FCS + 2% Penicillin-Streptomycin + 1% L-glutamine (Gibco)). For cell counting, lymphocytes were diluted 40x with Trypan Blue (ThermoFisher Scientific) and enumerated using a Neubauer ‘Improved’ haemocytometer (Biochrom). Alternatively, lymphocytes were counted using a MACSQuant flow cytometer (Miltenyi Biotec), using propidium iodide (PI) (Sigma Aldrich) or, in the case of hepatic lymphocytes, using CD45.2-Alexa647 (Biolegend) to distinguish between hepatocytes and lymphocytes, prior to PI administration and counting. Peripheral blood was acquired by tail vein puncture collected in Na^+^ heparin capillary tubes (Brand) and assayed in 96-well flat bottom plates (Corning). For CD8+ T cell stimulations, 2-3×10^6^ splenocytes or 1-2×10^5^ liver cells were incubated with SIINFEKL peptide (Peptides and Elephants, Henningsdorf) at a final concentration of 10μg/ml in the presence of Brefeldin A (eBioScience). Cells were incubated at 37°C, 5% CO_2_ for 5-6 hours, before incubation at 4°C overnight. For staining of cell surface markers and intracellular cytokines, cells were incubated for 1 hour at 4°C. Cells derived from the spleen or liver were fixed with 4% paraformaldehyde, and cells from peripheral blood were fixed with 1% paraformaldehyde between the extra- and intracellular staining steps. Data was acquired by flow cytometry using an LSRII or LSRFortessa (BD). Antibodies used for staining were as follows; BD: CD3 (500A2); eBioScience: CD8 (53-6.7), CD11a (M17/4), CD49d (R1-2), CD62L (MEL-14), CD44 (IM7), IFN-γ (XMG1.2), TNF-α (MP6-XT22) and IL-2 (JES6-5H4); ProImmune: H-2-K^b^-SIINFEKL pentamers.

### CFSE labelling of OT-I cells

Spleens from OT-I mice were lysed and cells washed twice in PBS without serum. Splenocytes resuspended at a density of 5×10^6^ cells/ml in PBS had 1:5,000 CFSE (ThermoFisher Scientific) added and were incubated in the dark at room temperature, with gentle inversion for 4 minutes. The labelling reaction was quenched with cRPMI and cells washed twice in cRPMI. Cells were recounted and 2×10^6^ cells were injected per mouse.

### Vaccination with OVA expressing recombinant adenovirus

To assess parasite liver load after vaccination with virus-expressed OVA, groups of C57BL/6 mice were immunized with recombinant human adenovirus serotype 5 (AdHu5) expressing full-length chicken ovalbumin^26^. Each mouse received 1×10^8^ infective units (ifu) in a volume of 100μl administered intramuscularly (50μl into each thigh). At the same time mice received OT-I splenocytes intravenously (2×10^6^ cells/mouse). 19 days after vaccination, vaccinated and naïve control mice were challenged with 10,000 WT, CSP^SIINFEKL^ or UIS4^SIINFEKL^ sporozoites administered intravenously. 42 hours after the challenge the livers were harvested and homogenised in Trizol (ThermoFisher Scientific) for total RNA isolation. Afterwards, cDNA was generated using the RETROScript Kit (Ambion). Quantitative real-time PCR was performed using the StepOnePlus Real-Time PCR System and Power SYBR Green PCR Master Mix (Applied Biosystems). Relative liver parasite levels were quantified using the ΔΔCt method comparing levels of *P. berghei 18S* rRNA normalised to mouse *GAPDH* mRNA^30^. To assess sterile protection, AdHu5 OVA-vaccinated and control mice received 2×10^6^ OT-I splenocytes one day prior to vaccination. 14 days later, all mice were challenged with 1,000 WT, CSP^SIINFEKL^ or UIS4^SIINFEKL^ sporozoites. Blood smears were taken from day 3-14 after challenge to determine the presence of blood stage parasites.

### Statistics

Data were analysed using FlowJo version 9.5.3 (Tree Star Inc., Oregon, USA), Microsoft Excel and GraphPad Prism v7 (GraphPad Software Inc., CA, USA). We used Mann-Whitney U test for analysing data that were not normally distributed and Welch’s t-test or one-way ANOVA with Tukey’s multiple comparison test for normally distributed data.

## Acknowledgements

S.J.D is a Jenner Investigator, Lister Institute Research Prize Fellow and Wellcome Trust Senior Fellow (106917/Z/15/Z). K.Matuschewski was supported by the Max Planck Society and grants from the European Commission (EviMalaR Network of Excellence #34) and the Chica and Heinz Schaller Foundation. O.S. was funded in part by the Laboratoire d’Excellence ParaFrap (ANR-11-LABX-0024). J.C.R.H. was funded by grants from The Royal Society (University Research Fellowship UF0762736/UF120026 and Project Grant RG130034) and the National Centre for the Replacement, Refinement & Reduction of Animals in Research (Project Grant NC/L000601/1). The funders had no role in study design, data collection and analysis, decision to publish, or preparation of the manuscript.

## Author contributions

K.Matuschewski, O.S. and J.C.R.H. designed the experiments; O.S. generated the transgenic parasites CSP^SIINFEKL^ and UIS4^SIINFEKL^; K.Müller, M.P.G., O.S. and J.C.R.H. performed experiments and analysed data; A.R.-S., A.V.S.H. and S.J.D. provided the adenovirus AdOVA; M.P.G. and J.C.R.H. wrote the paper; all authors commented and revised the manuscript.

## SUPPLEMENTARY EXPERIMENTAL PROCEDURES

**Generation of CSP^SIINFEKL^ and UIS4^SIINFEKL^ transgenic *P. berghei* parasite lines**

B3D-CSP^SIINFEKL^ plasmid was assembled by successive cloning of three fragments, CSP-C, CSP-B and CSP-A, obtained by PCR amplification from *P. berghei* ANKA genomic DNA followed by restriction enzyme digestion. These fragments correspond respectively to a 3’ homology region downstream of CSP (CSP-C, 0.7 kb), a fragment comprising the CSP ORF downstream of the SYIPSAEKI epitope followed by the CSP 3’ UTR (CSP-B, 0.8 kb) and a fragment comprising a 5’ promoter region followed by the CSP modified ORF where the SYIPSAEK coding sequence has been replaced by a SIINFEKL coding sequence (CSP-A, 1.8 kb). The resulting B3D-CSP^SIINFEKL^ plasmid, containing the *Toxoplasma gondii* dihydrofolate reductase/thymidylate synthase (*TgDHFR/TS*) pyrimethamine resistance cassette flanked by CSP-A and CSP-B on one side, and CSP-C on the other, was linearized with *Not*I and *Sac*II before transfection. Integration of the construct after double crossover homologous recombination results in replacement of the WT CSP gene by a modified copy containing the SIINFEKL coding sequence instead of the SYIPSAEKI coding sequence. The B3D-UIS4^SIINFEKL^ plasmid was assembled by successive cloning of three fragments, UIS4-A, UIS4-B and UIS4-C, obtained by PCR amplification from *P. berghei* ANKA genomic DNA followed by restriction enzyme digestion. These fragments correspond respectively to a fragment comprising a 5’ upstream sequence followed by the UIS4 entire ORF fused in frame to the SIINFEKL coding sequence (UIS4-A, 1.2 kb), to the UIS4 3’ UTR sequence (UIS4-B, 0.6 kb) and to a 3’ homology region downstream of UIS4 (UIS4-C, 0.9 kb). The resulting B3D-UIS4^SIINFEKL^ plasmid, containing the *TgDHFR/TS* pyrimethamine resistance cassette flanked by UIS4-A and UIS4-B on one side, and UIS4-C on the other, was linearized with *Sac*II and *Kpn*I before transfection. Integration of the construct after double crossover homologous recombination results in replacement of the WT UIS4 gene by a modified copy containing the SIINFEKL coding sequence just upstream of a STOP codon. *P. berghei* CSP^SIINFEKL^ and UIS4^SIINFEKL^ parasites were generated by transfection of *P. berghei* ANKA with linearized B3D-CSP^SIINFEKL^ and B3D-UIS4^SIINFEKL^ plasmids, respectively. Purified schizonts of WT *P. berghei* ANKA (clone c15cy1) were transfected with 5-10μg of linearized plasmid by electroporation using the AMAXA Nucleofector™ device (program U33), as described^39^, and immediately injected intravenously in the tail vein of a mouse. The day after transfection, pyrimethamine (70 mg/l) was administrated in the mouse drinking water, for selection of transgenic parasites. Transgenic clones were isolated after limiting dilution and injection into mice. Correct integration of the constructs and purity of the transgenic lines was verified by analytical PCR using primer combinations specific for the unmodified CSP or UIS4 locus, and for the 5’ and 3’ recombination events. All primers used in this study are indicated in Table S1.

## SUPPLEMENTARY FIGURE LEGENDS

**Suppl. Figure 1: Generation of transgenic CSP^SIINFEKL^ and UIS4^SIINFEKL^ *P. berghei* lines**

*Plasmodium berghei* parasites expressing the CD8+ T cell epitope of ovalbumin, SIINFEKL, in the context of CSP or UIS4 were generated using double homologous recombination, combining drug-resistance selection (through incorporation of the *dhfr/ts* gene from *Toxoplasma gondii*) and cloning by limiting dilution to select for correctly recombined parasites. **(a,b)** Diagrams illustrate the reverse genetics strategy. **(a)** In CSP^SIINFEKL^ SIINFEKL replaces the immunodominant CD8+ T cell epitope SYIPSAEK(I) of CSP. **(b)** In UIS4^SIINFEKL^ SIINFEKL is adjoined to the carboxyl-terminus of the UIS4 protein. Purified schizonts of WT *P. berghei* ANKA were transfected with linearized plasmid by electroporation as described^39^, and immediately injected intravenously in the tail vein of a mouse. The day after transfection, pyrimethamine (70 mg/l) was orally administered in the drinking water for selection of transgenic parasites. Transgenic clones were generated in mice by *in vivo* cloning by limiting dilution. Correct integration of the constructs and purity of the transgenic lines was verified by diagnostic PCR using primer combinations specific for the unmodified *CSP* or *UIS4* locus, and for the 5’ and 3’ recombination events as indicated by lines, arrows and expected fragment sizes. **(c)** Oocyst midgut infectivity of mosquitoes infected with WT, CSP^SIINFEKL^ or UIS4^SIINFEKL^. The mean percentage (±SD) of infected midguts was enumerated 10-14 days after infection (n= at least 7 infections). **(d)** Salivary glands were isolated from WT, CSP^SIINFEKL^ or UIS4^SIINFEKL^ infected mosquitoes and mean sporozoite numbers (±SD) were enumerated between 18-23 days after infection (n= at least 13 infections).

**Suppl. Figure 2: Sporozoite surface antigen induces a greater effector CD8+ T cell phenotype than EEF vacuolar membrane antigen.**

C57BL/6 mice (up to 5per group) received 2×10^6^ OT-I cells alone or were additionally immunised with 10,000 γ-radiation attenuated WT, CSP^SIINFEKL^ or UIS4^SIINFEKL^ sporozoites intravenously. Spleens and livers were harvested either 14 or 42 days later, and proportions of CD8+ T cells expressing effector surface markers were quantified. Flow cytometry plots show representative percentages of CD8+ T cells co-staining K^b^-SIINFEKL and markers of effector phenotype (CD11a^hi^, CD49d^hi^, CD62L^lo^, CD44^hi^).

**Suppl. Figure 3: Antigen experienced SIINFEKL-specific CD8+ T cells also produce TNF-α and IL-2.**

C57BL/6 mice (up to 5 per group) received 2×10^6^ OT-I cells alone or were additionally immunised with 10,000 γ-radiation attenuated WT, CSP^SIINFEKL^ or UIS4^SIINFEKL^ sporozoites intravenously. Spleens and livers were harvested either 14 or 42 days after immunisation and lymphocytes restimulated *ex vivo* with SIINFEKL peptide at 10μg/ml per well for 5-6 hours. The upper panel of bar charts show the percentage of CD11a+ TNF-α secreting CD8+ T cells, the bottom panel CD11a+ IL-2 secreting CD8+ T cells. This is a representation of one experiment from two experiments performed. Bar charts show mean values (±SEM) from representative experiments (*, p<0.05, **, p<0.01, ***, p<0.001; one-way ANOVA with Tukey’s multiple comparison test).

**Suppl. Table 1:**
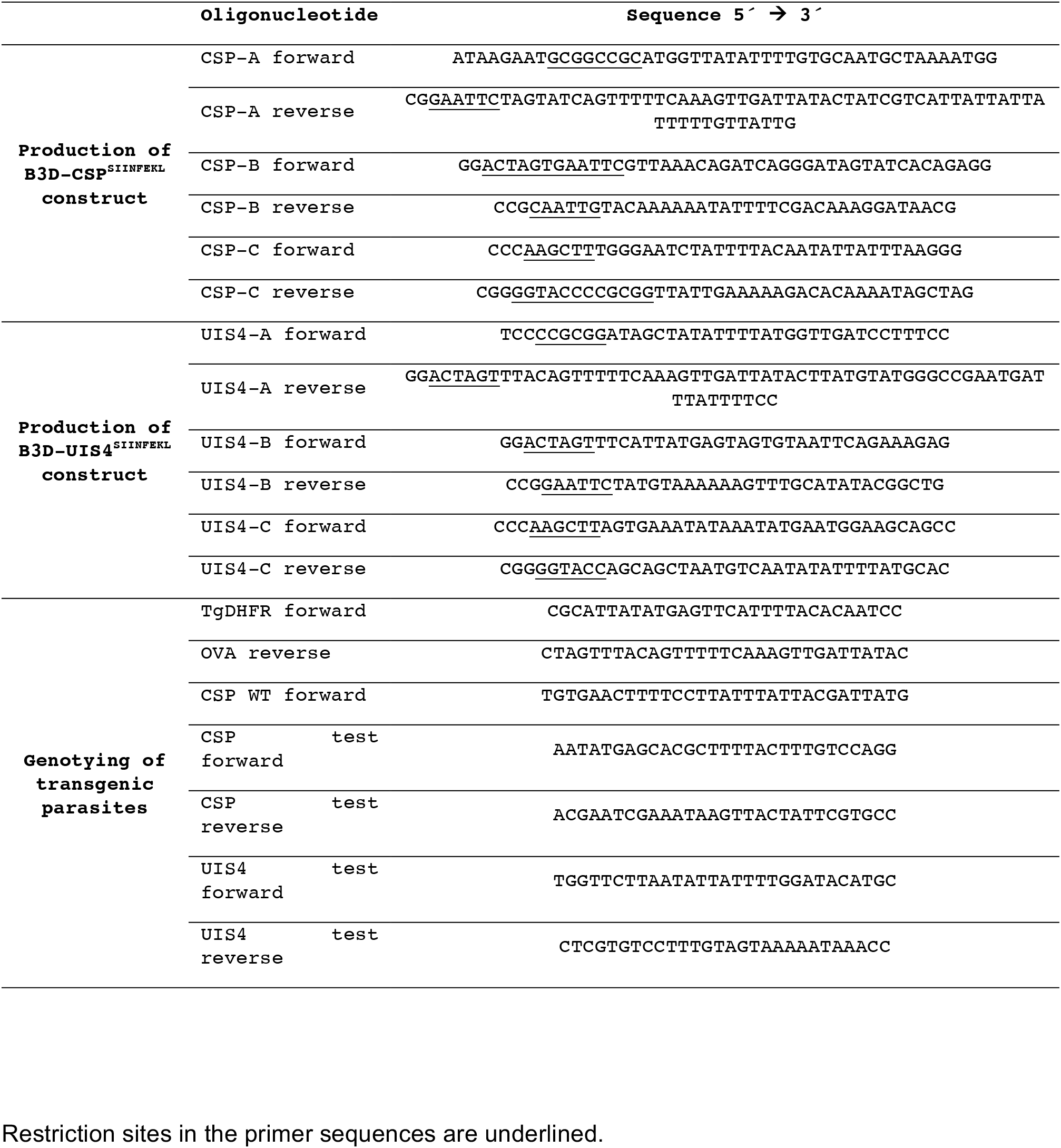
Primers used to generate plasmids and genotype parasites.

